# The respiratory phase causally modulates the readiness potential amplitude

**DOI:** 10.1101/2025.07.27.666697

**Authors:** Samuel J. Sandoval, Ye-Seung Suh, Kang-Yun Lee, Hyeong-Dong Park

## Abstract

Previous research has shown that the respiratory phase correlates with both voluntary action timing and readiness potential (RP) amplitude, but whether this relationship is causal or merely correlational remains unclear. Here, we experimentally manipulated breathing patterns to test if the respiratory phase causally influences the neural preparation for voluntary movement. Participants performed self-initiated button presses under four conditions: breathing in (BI), breathing out (BO), normal breathing (NB), and breath-holding (BH). Electroencephalographic recordings revealed that RP amplitude was significantly more negative during exhalation compared to inhalation and during breath-holding compared to normal breathing. These neural differences occurred without corresponding changes in behavioral measures (e.g., waiting times or retrospective timing judgments), indicating that respiratory effects on RP were not associated with altered action timing. These findings demonstrate that the respiratory phase causally modulates cortical motor preparation. We further propose that the brain optimizes voluntary action execution during respiratory phases when breathing-related motor activity is minimal, thereby revealing respiration as a fundamental organizing rhythm for voluntary behavior.

## Introduction

The readiness potential (RP), a negative cortical deflection preceding voluntary movements, has been central to debates about motor preparation and consciousness since its discovery by Kornhuber and Deecke (1965), particularly following Libet’s seminal experiments (1983). While traditionally interpreted as reflecting motor preparation, contemporary frameworks propose that the RP emerges from ongoing neural dynamics, either through stochastic fluctuations that reach a decision threshold (Schurger et al., 2012) or movements that coincide with specific phases of cortical potentials (Schmidt et al., 2016). Despite recent mechanistic insights using neuronal recordings, the RP’s fundamental nature remains contested (Gavenas et al., 2024; Schurger et al., 2021). If the RP reflects spontaneous fluctuations rather than dedicated motor preparation, what factors could shape these fluctuations? Emerging evidence suggests that respiration, increasingly recognized as rhythmically organizing perception, cognition, and behavior (Heck et al., 2019; Johannknecht & Kayser, 2022; Kluger et al., 2021, 2023; Perl et al., 2019), could potentially influence these neural dynamics. Previously, our group reported that the respiratory phase is associated with both voluntary action timing and RP amplitude (Park et al., 2020, 2022). Using classical Kornhuber and Libet tasks, it was found that participants unconsciously preferred to initiate self-initiated movements during the exhalation phase, despite having complete temporal freedom. Moreover, RP amplitude varied systematically with respiratory phase in each trial. These findings were subsequently extended to mental imagery tasks, suggesting that respiratory influences on action preparation extend beyond overt movement (Park et al., 2022). In these studies, we further proposed that such coupling might serve to minimize potential motor competition between respiratory commands and voluntary movement signals at either cortical or subcortical levels. According to this motor competition hypothesis, participants might unconsciously initiate voluntary actions during exhalation (i.e., the “passive” phase of breathing) to avoid interference between concurrent motor processes.

The present study aims to address a key follow-up question: Does the respiratory phase causally impact the amplitude of RP? Initial observations remain fundamentally correlational. While the behavioral coupling between respiration and voluntary action timing is well-established, the relationship between respiratory phase and RP amplitude requires further investigation (Schurger et al., 2021). The observed correlation between RP and respiration could reflect two possibilities: either respiratory phase causally influences the neural preparation for movement, or this correlation is merely a secondary consequence of the established behavioral tendency for movements to occur during specific respiratory phases. In other words, if movements preferentially occur during exhalation, and the RP always precedes movement, then RP measurements could be biased toward the exhalation phase without any direct causal relationship between breathing and neural preparation. The present study directly addresses this limitation by experimentally manipulating respiratory patterns to test for causal influence on RP amplitude. If respiration plays a causal role in shaping the neural dynamics underlying voluntary action, then controlled alterations in breathing patterns during a voluntary action task should produce corresponding changes in RP characteristics. Based on our previous findings and the motor competition hypothesis, we hypothesized that RP amplitude would be inhibited during breathing in (BI) compared to breathing out (BO) conditions. In addition, we conducted an exploratory comparison of RP amplitude between normal breathing (NB) and breath-holding (BH) conditions, where the generation of a new breathing cycle is absent.

## Methods

### Physiological Data Acquisition

Continuous EEG data were collected from 20 scalp locations using the CGX Quick-20 dry electrode system at a sampling rate of 500 Hz. We used A1 (left ear) as reference, with A2 as the complementary reference electrode. All EEG signals were displayed in real-time via BrainVision Recorder for CGX software. Concurrent physiological signals were recorded using the CGX AIM physiological monitor at a sampling rate of 500 Hz. Electrocardiogram (ECG) signals were obtained using bipolar electrodes positioned below the collarbones in the second intercostal space along the midclavicular line on both sides. The same ECG electrodes were also used to collect respiratory signals through impedance pneumography: a small alternating current passed between the chest electrodes, and changes in thoracic impedance caused by lung inflation/deflation were measured to track breathing. In both experiments, stimulus presentation was controlled by a dedicated computer running PsychoPy (v2022.2.5). Trigger signals were simultaneously sent to recording computers by a StimTrigger (Cedrus/CGX) using 16-bit wireless transmission, allowing precise temporal synchronization between stimulus events and physiological recordings.

### Participants

Thirty healthy volunteers (11 females, 2 ambidextrous; mean age: 26.33 ± 4.22 years) participated in the study. All participants reported no history of cardiovascular disease, neurological or psychiatric disorders. One participant was excluded from the analysis due to excessive movement artifacts. Two participants were further excluded due to an issue (i.e., age) related to the IRB. The data from 27 subjects were analyzed. All participants provided written informed consent before the experiment and received monetary compensation for their participation. The study was approved by the local ethics committee (TMU-Joint Institutional Review Board) and conducted in accordance with the Declaration of Helsinki.

### Experimental design

The experiment employed a modified Libet paradigm to investigate the influence of differential respiratory patterns on breathing-action coupling and the RP. We designed four distinct experimental conditions, each manipulating the participant’s breathing pattern in relation to voluntary action: NB, BH, BO, and BI. During the NB condition, participants were instructed to “breathe naturally and normally as they would during daily activities,” and to initiate button presses whenever they spontaneously felt the urge to do so. They were explicitly asked not to deliberately alter their breathing pattern, avoid deep or controlled breaths, and to focus instead on naturally and spontaneously deciding the moment of action initiation. This baseline condition captured spontaneous action timing without any respiratory manipulation. During the BH condition, participants were explicitly instructed to inhale deeply upon seeing a cue (a brief visual circle displayed at the center of the screen, lasting 3 seconds) and subsequently hold their breath while initiating the voluntary button press. They were permitted to initiate the button press at any time during the breath-holding period, but were clearly instructed to maintain the breath hold until the action was initiated. Participants resumed normal breathing immediately after the action was executed. During the BO condition, participants received explicit instructions to initiate their voluntary button presses exclusively during the exhalation phase. During the BI condition, participants were instructed to initiate the button press while breathing in. Importantly, during the BO and BI conditions, participants were instructed to breathe naturally and normally without deliberately modulating their breathing patterns.

### Procedure

Participants performed the task while seated 60 cm from a monitor displaying an analogue clock (circular outline, 2° visual angle radius) with 12 major and 60 minor tick marks. Each trial began with a green pointer (0.4° diameter) appearing at a random position and rotating clockwise at a reduced speed (i.e., 10.24 seconds per rotation, approximately four times slower than the traditional Libet paradigm). This rotation period was deliberately chosen to span approximately two complete respiratory cycles, ensuring participants could complete at least one full breath before reaching the decision point, regardless of which phase of breathing they were in, when the trial began. This modification follows prior studies that lengthened the Libet clock for investigating non-motoric influences on the RP (Alexander et al., 2016; Park et al., 2022). It provided sufficient time for natural breathing and cognitive processing before movement initiation, which was essential for testing our hypotheses regarding respiratory manipulation. Participants were told to wait until the clock hand completed at least one quarter rotation (i.e., 2.55 seconds) before initiating action. To facilitate task comprehension and minimize cognitive load, we included two visual markers (semi-transparent black lines, 0.5 alpha, 2.0-pixel width) extending from the center to the initial dot position on the clock face indicating the starting angle (θ0) and the position marking one quarter of a rotation (θ0 + π/2), helping participants identify when they could initiate their response. Instructions emphasized spontaneous timing choices, explicitly discouraging pre-planning or regular interval pressing. After action initiation, the clock hand disappeared for 4 seconds before reappearing at the action position to facilitate retrospective reporting of intention timing (i.e., W-time). Between trials, a variable interval (i.e., 4-8 seconds) was interleaved, allowing both neural activities and respiratory patterns to return to baseline, ensuring comparable physiological states at each trial onset.

A critical aspect of our design was ensuring that participants maintained as natural a breathing pattern as possible, even during the manipulation conditions. We specifically instructed participants not to exaggerate any respiratory phase or significantly alter their normal breathing rhythm. For the BH condition, we asked participants to take a natural deep breath before holding, rather than a forced maximal inhalation. For the BO and BI conditions, we emphasized that participants should maintain their typical breathing rate and depth, only adjusting the timing of their voluntary action to coincide with the required phase. Visual animations at the beginning of each block reinforced these instructions.

The experiment consisted of 12 blocks (three per condition, 25 trials each, totaling 75 trials per condition) presented in randomized order. Each block was followed by a mandatory 60-second break, with additional rest time available if needed. Before the main experiment, participants completed a training session to ensure familiarity with all breathing conditions and timing estimation procedures. The complete experimental session, including setup and training, lasted approximately 2.5 hours.

### Behavioral Analysis

Behavioral data were analyzed using Python (v3.11.2) to calculate waiting times (i.e., from stimulus onset to button press) and W-times (i.e., retrospectively reported awareness of intention). Given the 10.24 clock rotation, we converted reported angles to time (degrees × 10.24/360). To account for the circular clock face, we computed the minimum angular distance between the reported intention and the actual button press positions, ensuring an accurate measurement of the intention-action latency. Data was averaged per participant and condition. We examined whether respiratory conditions influenced voluntary action timing through two planned comparisons: BI vs. BO and NB vs. BH. Given the non-normal distribution typical of voluntary action paradigms, we employed the Wilcoxon signed-rank test (Wilcoxon, 1945), a non-parametric alternative appropriate for our within-subjects design. Effect sizes were quantified using Cohen’s d for paired samples. To complement our analysis, we calculated Bayes Factors (BF10) using the Jeffrey-Zellner-Siow prior (Rouder et al., 2009), which quantifies relative evidence for the alternative hypothesis (conditions differ) versus the null (no difference). We interpreted BF10 following standard conventions: < 1/3 (moderate evidence for H0), 1/3–3 (anecdotal evidence), 3–10 (moderate evidence for H1), and > 10 (strong evidence for H1).

### Respiratory Signal Analysis

We analyzed respiratory signals to determine whether voluntary actions were systematically synchronized with specific phases of the breathing cycle. Respiration data were processed using the Fieldtrip toolbox (Oostenveld et al., 2011) in MATLAB (R2022b). For preprocessing, respiratory signals were bandpass filtered (0.2-0.8 Hz, second-order Butterworth) to isolate natural breathing rhythms (typically 12-30 breaths per minute) while eliminating artifacts and signal drift. We then applied the Hilbert transform to extract instantaneous phase information, yielding a continuous measure (0-2π) that identified each moment in the respiratory cycle, with 0-π representing exhalation and π-2π representing inhalation. Then we extracted the respiratory phase at the timing of voluntary action onset in each trial.

### EEG Analysis

All preprocessing was conducted using the Fieldtrip toolbox (Oostenveld et al., 2011). EEG data were offline filtered between 0.1 and 10 Hz. Independent component analysis was conducted, and stereotypical independent components reflecting eye movements and eye blinks were excluded (Delorme & Makeig, 2004). The RP was computed from EEG signals locked to the button-pressed time, within a 5-second time window (-4 to 1 second, relative to the markers). Baseline correction was applied using a time window of [-4 to -3] seconds. Trials showing excessive noise (> 3 SD) were excluded from further analysis. Because RPs are known to be strongest at the central region, we report RP results over the Cz electrode following previous RP studies (Maoz et al., 2019; Schultze-Kraft et al., 2016).

The significance of the difference in RP between two planned comparisons (i.e., BI vs. BO; NB vs. BH) was tested using the cluster-based permutation test in the temporal dimension over the [-1 to 0] seconds time window, as implemented in the FieldTrip toolbox (Oostenveld et al., 2011). This procedure extracts temporal clusters that show significant differences between conditions without any a priori specification of time windows and intrinsically corrects for multiple comparisons in time. The permutation p-value corresponds to the proportion of shuffled maximal cluster-level statistics (1000 times) that exceeds the observed original cluster-level test statistics.

## Results

### Behavioral Performance

To test whether experimentally altered respiratory phases causally influence the RP amplitude, we conducted two planned comparisons: BI vs. BO and NB vs. BH. Before comparing RP amplitude between these conditions, we checked the behavioral performance (e.g., waiting time and W-time). The overall mean waiting time across all conditions was 6.17 ± 0.19 seconds (mean ± SE throughout), consistent with our previous study using extended Libet clock paradigms (6.30 ± 0.66 seconds in Park et al. (2022)). The distribution of waiting times showed the characteristic right-skewed shape typical of voluntary action tasks. Participants’ retrospective reports of their conscious intention to move (W-time) occurred on average -244 ± 47 milliseconds before the actual button press, aligning with typical findings in voluntary action studies where conscious awareness of intention precedes movement by approximately 200-300 milliseconds (Libet et al., 1983; Park et al., 2020, 2022).

Analysis of waiting times revealed no significant differences between breathing conditions. In the BI vs. BO comparison, participants demonstrated similar waiting times (BI: 6.36 ± 0.17 s; BO: 6.37 ± 0.18 s; Fig. 1a). The Wilcoxon signed-rank test confirmed this similarity (p = 0.750), with a negligible effect size (Cohen’s d = -0.04) and a Bayes Factor of 1.02 providing no evidence in favour of either hypothesis. The retrospective timing judgments (W-times) showed a comparable pattern, with no significant difference between conditions (BI: -0.259 ± 0.046 s; BO: -0.240 ± 0.046 s; p = 0.313; Cohen’s d = -0.206; BF₁₀ = 1.69).

**Figure 1.**
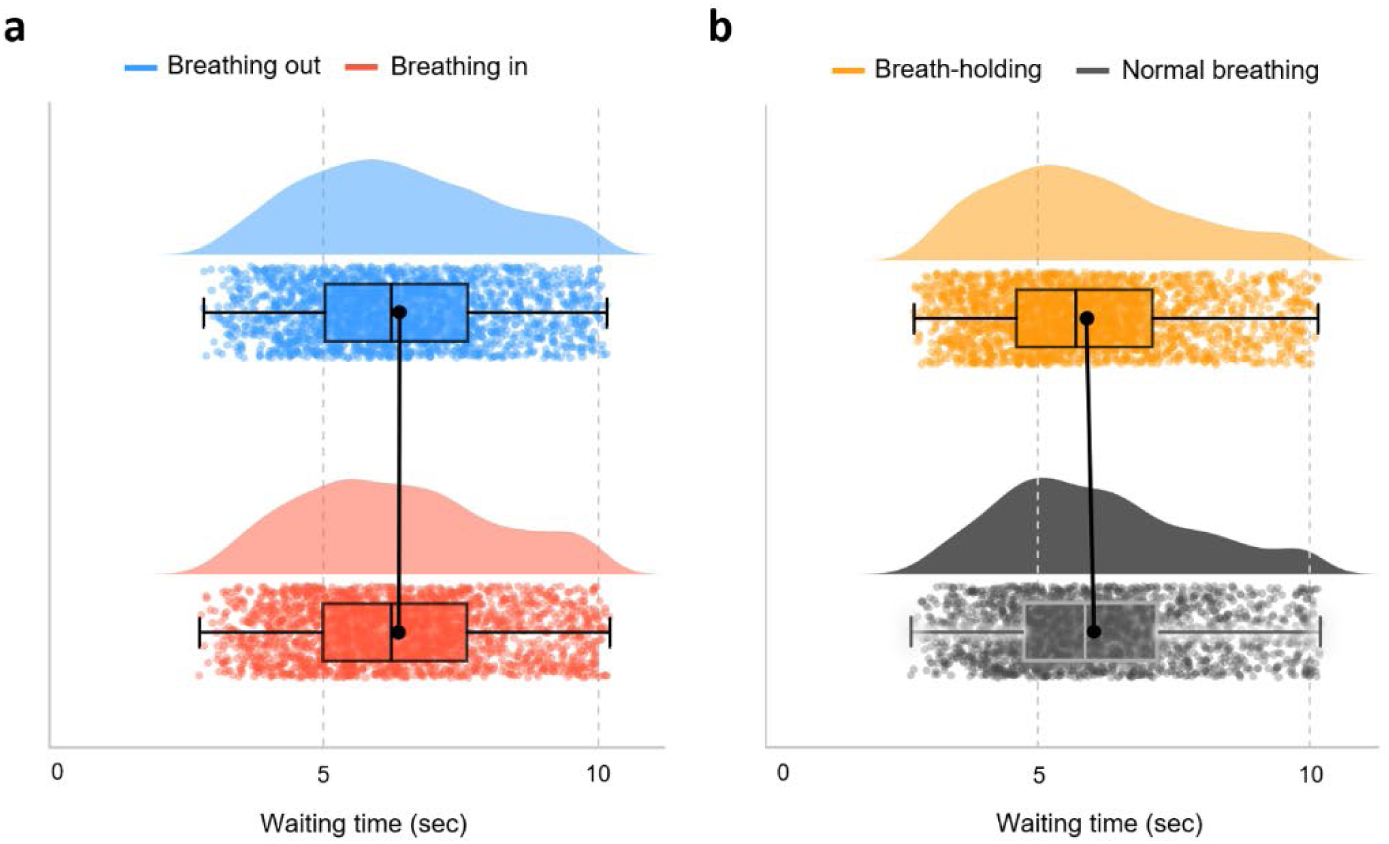
Distribution of waiting times across respiratory conditions. Raincloud plots showing waiting time distributions for **a.** breathing in (BI) vs. breathing out (BO) conditions, and **b.** normal breathing (NB) vs. breath-holding (BH) conditions. Each plot displays the probability density (half-violin plot), individual trials (dots), and summary statistics (box plot with median line and IQR). The bold black line connects the condition means. All distributions show the characteristic right-skewed pattern typical of voluntary action timing tasks. For the BI versus BO comparison, waiting times were nearly identical (BI: 6.36 ± 0.17 s; BO: 6.37 ± 0.18 s; p = 0.750, Cohen’s d = -0.04, BF₁₀ = 1.02). Similarly, the NB versus BH comparison revealed no significant differences (NB: 6.03 ± 0.19 s; BH: 5.90 ± 0.21 s; p = 0.170, Cohen’s d = 0.27, BF₁₀ = 2.34).

The comparison between NB and BH conditions yielded similar results. Waiting times did not differ significantly between conditions (NB: 6.03 ± 0.19 s; BH: 5.90 ± 0.21 s; p = 0.170; Cohen’s d = 0.27; Fig. 1b), with Bayesian analysis providing only anecdotal evidence (BF₁₀ = 2.34) that was insufficient to support either the null or alternative hypothesis. W-time measurements likewise showed no significant difference (NB: -0.225 ± 0.049 s; BH: -0.252 ± 0.047 s; p = 0.202; Cohen’s d = 0.297; BF₁₀ = 2.85), indicating consistent timing of conscious intention across respiratory states.

Collectively, these behavioural findings demonstrate that participants successfully maintained voluntary action timing within experimental constraints regardless of respiratory manipulation. The stability of both waiting times and W-times across all conditions suggests that experimental manipulation of respiratory phases in our task design did not induce altered temporal dynamics of voluntary action initiation or conscious awareness across conditions.

### EEG results

We tested the idea that the RP amplitude depends on the experimentally modulated respiratory phase. We first compared the RP amplitude between the BI and BO conditions. For this, we excluded trials that were not under the designated respiratory phase criterion in both conditions (BO: 0 to 180 degrees; BI: 180 to 360 degrees; see Fig. 2a). After trial exclusion, 57.0 ± 2.04 and 50.4 ± 2.11 epochs were averaged to compute RP for BO and BI conditions, respectively. RP amplitude significantly differed between BI and BO conditions over the electrode Cz (cluster-based permutation test in temporal dimension; permutation p = 0.022) between -520 and 0 ms (Fig. 2b). RP amplitude was more prominent (i.e., negative) in BO compared to BI, supporting the proposed motor competition hypothesis (see Discussion). Then, we further applied a shifted criterion for the trial exclusion (BO: 30 to 210 degrees; BI: 210 to 30 degrees; see Supplementary Fig. 1a) to maximize the total number of trials and to preserve participants’ natural preference during the task. After trial exclusion with shifted criterion, 55.5 ± 2.29 and 55.9 ± 2.00 epochs were averaged to compute RP for BO and BI conditions, respectively. Based on this shifted exclusion criterion, we also found similar significant RP differences between BO and BI conditions (Supplementary Fig. 1b).

**Figure 2.**
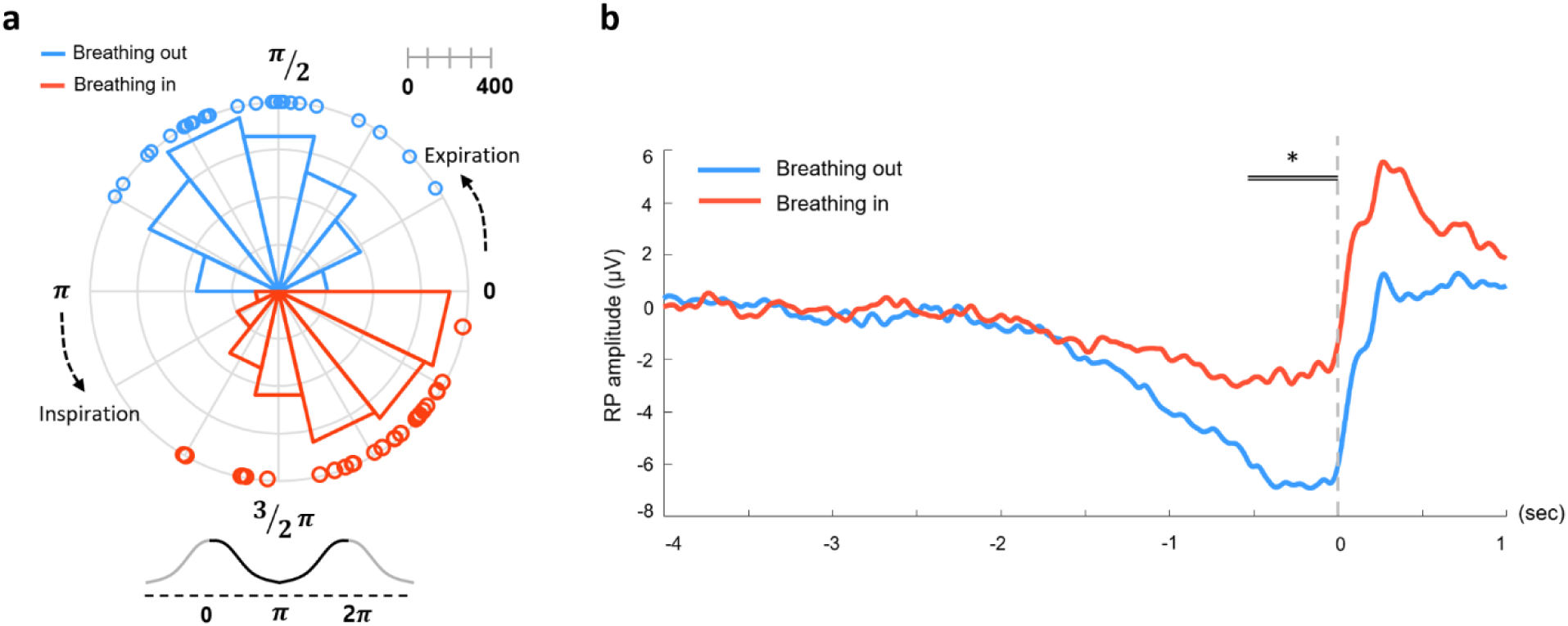
Respiratory phases and RP amplitudes in breathing in (BI) and breathing out (BO) conditions. **a.** Distribution of respiratory phases at the timing of button presses in BI (red) and BO (blue) conditions (N = 27). The polar histogram shows the distribution of all the button presses from 27 participants. Empty circles represent each participant’s mean respiration phase at button press. The scale bar on the top-right indicates the number of action onsets in the polar histogram. **b.** The RP waveforms during BI (red) and BO (blue) conditions. RP during BO was significantly more negative compared to the BI condition (permutation *p* = 0.022). [-4 to -3 s] time window was set as a baseline period. The black double lines above indicate the time window in which significant RP was observed (-520 ∼ 0 ms).

Next, we investigated whether RP amplitudes differed between the NB and BH conditions. During the NB condition (Fig. 3a, black), participants preferred to initiate voluntary action during exhalation rather than inhalation, replicating previous observations. During the BH condition (Fig. 3a, orange), the respiratory phases were more frequently observed in the inhalation phase than in the exhalation phase, presumably reflecting a slow and slight increase in tidal volume during breath-holding. The RP amplitude significantly differed between NB and BH conditions over the electrode Cz (cluster-based permutation test in temporal dimension; permutation p = 0.026) between -272 and 0 ms (Fig. 3b). RP amplitude was more prominent (i.e., negative) in BH compared to NB.

**Figure 3.**
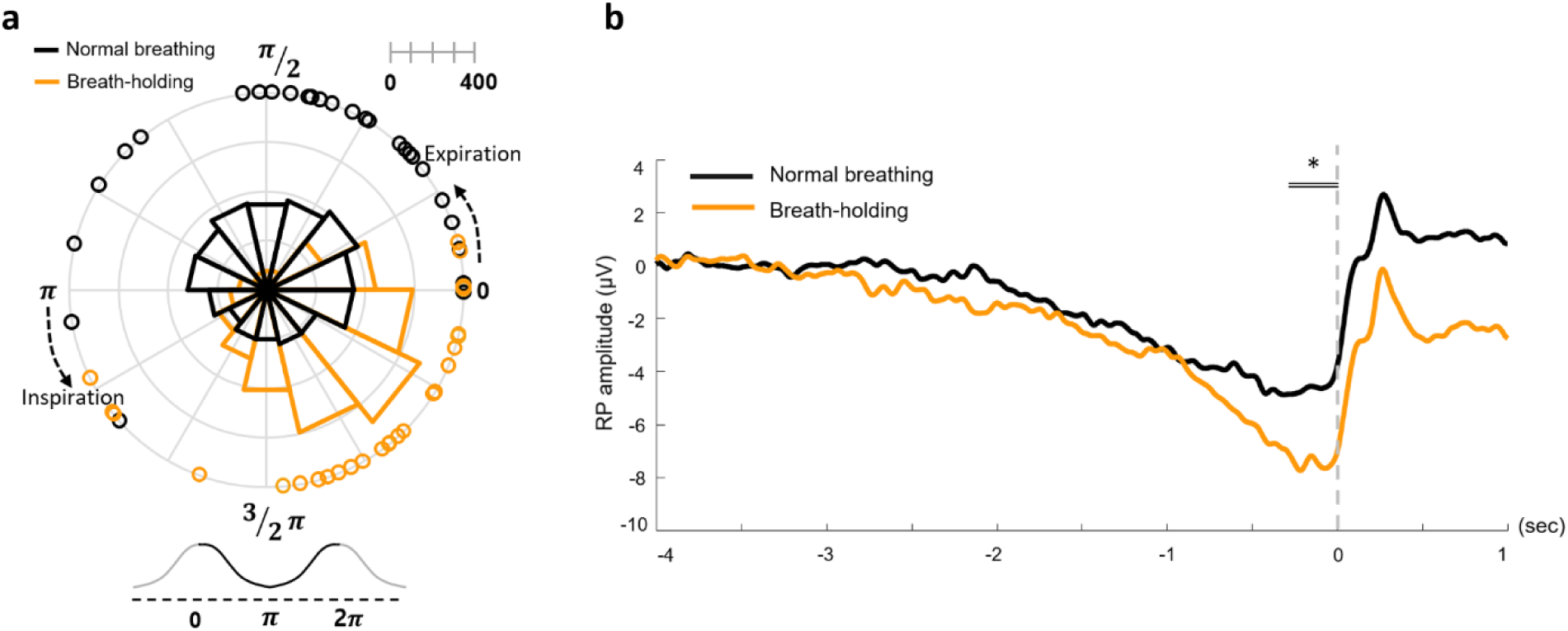
Respiratory phases and RP amplitudes in normal breathing (NB) and breath-holding (BH) conditions. **a.** Distribution of respiratory phases at the timing of button presses in NB (black) and BH (orange) conditions (N = 27). The polar histogram shows the distribution of all the button presses from 27 participants. Empty circles represent each participant’s mean respiration phase at button press. The scale bar on the top-right indicates the number of action onsets in the polar histogram. **b.** The RP waveforms during NB (black) and BH (orange) conditions. RP during BH was significantly more negative compared to the NB condition (permutation *p* = 0.026). [-4 to -3 s] time window was set as a baseline period. The black double lines above indicate the time window in which significant RP was observed (-272 ∼ 0 ms).

## Discussion

The present study investigated the causal influence of differential respiratory patterns on the neural preparation for voluntary action, as indexed by the RP. Our findings revealed robust neural differences between respiratory conditions (i.e., BI vs. BO, and NB vs. BH). Specifically, both comparisons showed differential RP amplitude during the final 500 milliseconds before movement execution. This temporal specificity suggests that respiratory state influences the immediate preparatory processes for motor execution, so-called the late RP (Schurger et al., 2021). Neither the presence versus absence of respiratory rhythm nor the coupling of actions to specific respiratory phases significantly altered waiting times or retrospective reports, excluding the possibility that observed RP differences were associated with differential behavioral performance.

### From correlation to causation

We previously reported that correlational relationships exist between the timing of voluntary action, RP amplitude, and breathing phase (Park et al., 2020, 2022). The correlation between RP and voluntary action was well established as early as the 1960s (Kornhuber & Deecke, 1965), although the causal nature of this relationship remains unknown. In our earlier work, we further demonstrated that the respiratory phase is also correlated with both the timing of voluntary action and the amplitude of the RP. More specifically, for the correlation between RP and respiratory phase, we employed a within-trial analysis to test whether respiratory phase is correlated with the RP waveform in each trial. Recently, Schurger et al. (2021) noted that our observation could reflect an incidental relationship rather than a causal one, as the correlation between RP and respiratory phase may be confounded by the other correlation between respiration and voluntary action. In other words, the observed correlation between the RP and respiration could be driven by the fact that voluntary action is time-locked to the expiratory phase. Thus, in the current study, we designed a follow-up experiment to test whether a causal relationship exists between the RP and the respiratory phase. For this, we experimentally manipulated the respiratory phase by instructing participants to initiate button presses under differential respiratory conditions. Then, we applied a between-trial analysis to reveal how differential respiratory phase causally impacts the RP amplitude. If the respiratory phase had merely correlated with RP through its association with action timing, we would have observed similar RP amplitudes across all respiratory conditions. Instead, we found significant differences in RP amplitude between respiratory states in both planned comparisons, suggesting that the respiratory phase has a causal impact on RP amplitudes.

### Potential underlying mechanisms

Importantly, we observed that the RP amplitude was enhanced (i.e., more negative) during the expiration compared to the inhalation period. This finding is in accordance with the explanation we previously proposed, the so-called motor competition hypothesis (Park et al., 2020), which states that the involuntary breathing-related and the voluntary limb-related motor commands compete for shared motor resources. It is well known that the pre-Bötzinger complex, located in the ventral medulla oblongata, generates the inspiratory phase of each breathing cycle. Most breathing-related pre-Bötzinger complex neurons exhibit peak activity during inspiration and markedly reduced activity during expiration (Del Negro et al., 2018; Menuet et al., 2020). According to this idea, the unconscious preference for initiating voluntary movements during expiration reflects optimal timing when respiratory motor systems are least active. Thus, the motor competition hypothesis predicts that RP amplitude would be reduced during inhalation, resulting from the actual occurrence of motor competition. In contrast, the RP amplitude would be enhanced or intact during exhalation, as there would be minimal competition between the two commands. We also observed more negative RP during the BH condition compared to the NB condition. Although we had a firm prediction on the comparison between BI and BO conditions, we predicted that two opposite results could be observed for the comparison between the BH and NB conditions. The first possibility, consistent with the motor competition hypothesis, suggests that eliminating respiratory rhythm generation during breath-holding would remove a source of neural interference, allowing motor preparatory signals to develop without competition. The alternative possibility is that the continuous motor effort required to maintain breath-holding might itself compete with voluntary movement preparation, potentially reducing RP amplitude, in which case we would have observed decreased RP during the BH condition compared to the NB condition. Our results support the first possibility. Taken together, our findings provide neural evidence in support of the motor competition hypothesis.

Notably, several non-mutually exclusive mechanisms may further explain our results. First, from a biomechanical perspective, exhalation stabilizes the trunk musculature through increased activation of abdominal muscles and a concurrent rise in intra-abdominal pressure, potentially creating an optimal platform for action execution (Hodges & Gandevia, 2000). This biomechanical state is ideal for executing movements that require postural control or force transmission, such as punching or precise motor tasks. Together with previous evidence showing that voluntary actions tend to align with exhalation, our results support the idea that the brain may utilize this stable respiratory phase to optimize motor execution. Second, the respiratory cycle may orchestrate a natural perception-to-action sequence, where inhalation enhances sensory sampling and cognitive processing in parieto-occipital areas (Kluger et al., 2021; Perl et al., 2019), while exhalation may provide the optimal window for motor execution over fronto-central regions, as observed in the present study. Recent findings by Kluger et al. (2023) demonstrated that respiration modulates the aperiodic component of neural activity (1/f slope), reflecting shifts in excitation-inhibition balance across the respiratory cycle. On the other hand, according to the leaky stochastic accumulator model proposed by Schurger et al. (2012), the RP does not represent a specific decision signal but rather reflects ongoing stochastic fluctuations in neural activity that occasionally cross a threshold for movement initiation. Our results are compatible with these frameworks: respiratory states could systematically bias these stochastic fluctuations, making threshold crossing more probable during exhalation when the excitation-inhibition balance favors motor output. The enhanced RP amplitude we observed during exhalation may thus reflect respiratory-induced shifts in the baseline of neural fluctuations rather than changes in a dedicated preparatory signal per se. These observations also align with a recent theoretical proposal by Mori & Sakano (2024), who suggested that one respiratory cycle constitutes a fundamental unit of behavioral decision-making in mammals. In their model, inhalation serves as a window for bottom-up sensory sampling, while exhalation provides a privileged phase for top-down decision and motor output. This cyclical organization of cognition and action offers a compelling temporal framework for voluntary behavior, consistent with the idea that the motor system may entrain itself to the exhalation phase, thereby reducing motor competition and optimizing both physiological stability and neural dynamics.

## Conclusion

The present study advances our understanding of voluntary action by establishing a causal link between respiratory state and cortical motor preparation, as indexed by the RP. By experimentally manipulating respiratory conditions, we demonstrated that the breathing phase directly modulates RP amplitude, with enhanced motor readiness during exhalation and breath-holding compared to inhalation and normal breathing. These findings could potentially advance our understanding of the principles behind sensorimotor integration and action execution. Central to this understanding is the motor competition hypothesis, which provides a potential mechanistic framework for how respiratory and voluntary motor systems could interact at the neural level. This competition-based mechanism is further supported by converging evidence from biomechanical optimization and neural excitability perspectives, suggesting that respiration serves as a fundamental organizing rhythm for voluntary behavior. Future investigations should explore whether this competitive interaction extends to more complex motor behaviors and decision-making processes, and whether individual differences in respiratory-motor coupling relate to motor performance or neurological differences in both healthy and clinical populations.

## Acknowledgements

This work was supported by the National Research Foundation of Korea (NRF) grant funded by the Korea government (MSIT)(RS-2024-00337437) to HDP.

## Data Availability Statement

The data supporting the findings of this study will be made publicly available in a suitable public repository upon acceptance of the manuscript. The accession link or DOI will be provided in the final published version.

