## Supplementary Figure 1 for "The respiratory phase causally modulates the readiness potential amplitude"

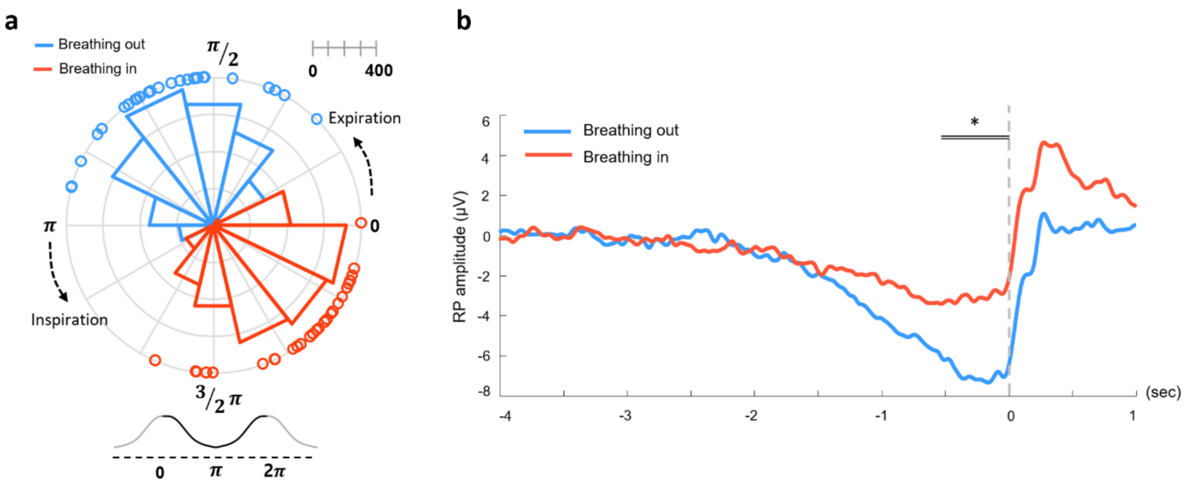


**Supplementary Figure 1.** Respiratory phases and RP amplitudes in the breathing in (BI) and breathing out (BO) conditions with a shifted criteria for trial exclusion. **a.** Distribution of respiratory phases at the timing of button presses in BI (red) and BO (blue) conditions (N = 27). The polar histogram shows the distribution of all the button presses from 27 participants. Empty circles represent each participant's mean respiration phase at button press. The scale bar on the top-right indicates the number of action onsets in the polar histogram. **b.** The RP waveforms during BI (red) and BO (blue) conditions. RP during BO was significantly more negative compared to the BI condition (permutation p = 0.018). [-4 to -3 s] time window was set as a baseline period. The black double lines above indicate the time window that significant RP was observed (-520 ~ 0 ms).
